# Fossils know it best: using a new set of fossil calibrations to improve the temporal phylogenetic framework of murid rodents (Rodentia: Myomorpha: Muroidea: Muridae)

**DOI:** 10.1101/180398

**Authors:** Tatiana Aghová, Yuri Kimura, Josef Bryja, Gauthier Dobigny, Laurent Granjon, Gael J. Kergoat

## Abstract

Murid rodents (Rodentia: Myomorpha: Muroidea: Muridae) represent the most diverse and abundant mammalian group. In this study, we reconstruct a dated phylogeny of the family using a multilocus dataset (six nuclear and nine mitochondrial gene fragments) encompassing 160 species representing 82 distinct murid genera from four extant subfamilies (Deomyinae, Gerbillinae, Lophiomyinae, and Murinae). In comparison with previous studies on murid or muroid rodents, our work stands out for the implementation of multiple fossil constraints within the Muridae thanks to a thorough review of the fossil record. Before being assigned to specific nodes of the phylogeny, all potential fossil constraints were carefully assessed; they were also subjected to several cross-validation analyses. The resulting phylogeny is consistent with previous phylogenetic studies on murids, and recovers the monophyly of all sampled murid subfamilies and tribes. Based on nine controlled fossil calibrations, our inferred temporal timeframe indicates that the murid family likely originated in the course of the Early Miocene, 23.0-16.0 million years ago (Ma), and that most major lineages (i.e. tribes) have started diversifying *ca.* 10 Ma. Historical biogeography analyses support the Paleotropical origin for the family, with an initial internal split (vicariance event) followed by subsequent migrations between Afrotropical and Indomalayan lineages. During the course of their diversification, the biogeographic pattern of murids is marked by several dispersal events toward the Australasian and the Palearctic regions, mostly from the Indomalaya. The Afrotropical region was also secondarily colonized at least three times from the Indomalaya, indicating that the latter region has acted as a major centre of diversification for the family.

## 1. Introduction

With about 150 genera and more than 730 recognized species, Muridae is the most diverse family of mammals (Musser and Carleton, 2005). Collectively murids have colonized highly distinct ecological niches, adapting to a wide array of environments ranging from warm (deserts or tropical forests) to cold habitats (high altitude mountain ranges, tundra; Vaughan et al., 2011). Life habits in murids are also diverse, as the family encompasses amphibious, arboreal, fossorial, or terrestrial taxa (Michaux et al., 2007; Musser and Carleton, 2005).

All murid species are native to the Old World (Musser and Carleton, 2005), but some species (especially the black rat *Rattus rattus* Linnaeus, the Norway rat *Rattus norvegicus* (Berkenhout) and the house mouse *Mus musculus* Linnaeus) now have a worldwide distribution due to commensalism and dissemination by humans. Murid species diversity is especially high in the Australasian and Indomalayan regions which accommodate half of the species diversity of the family (Rowe et al., 2016a). Second to that is the species diversity in the Afrotropical region (more than 200 species; Musser and Carleton, 2005). By contrast, there are much less native murid taxa in the Palearctic region (e.g. *Apodemus* Kaup, *Diplothrix* Thomas, or *Tokudaia* Kuroda).

The history of murid systematics is complex and convoluted with numerous changes occurring in the past sixty years (see Table 1 for a summary). Simpson (1945) divided representatives of family Muridae (as currently understood) into two separate families: Cricetidae (with subfamilies Gerbillinae, Lophiomyinae and others) and Muridae (subfamilies Murinae and Otomyinae). Chaline et al. (1977) considered “murid” rodents to belong to four families: Cricetidae (including Lophiomyinae), Gerbillidae, Muridae (exclusively Murinae) and Nesomyidae (including Otomyinae). Lavocat (1978) simplified this classification by recognizing only two families: Muridae (Murinae) and Nesomyidae (in which he included Gerbillinae, Lophiomyinae and Otomyinae). Another major change was later made by Carleton and Musser (1984), who defined family Muridae in the broad sense with no less than 14 subfamilies (including Gerbillinae, Lophiomyinae, Murinae and Otomyinae). Following the introduction of molecular systematics, changes in the classification of family Muridae continued at a fast rate. Using molecular phylogenetics Chevret et al. (1993a) demonstrated that *Acomys* I. Geoffroy is not a member of the subfamily Murinae but belongs to a separate monophyletic clade including *Deomys* Thomas, *Lophuromys* Peters and *Uranomys* Dollman. All four genera were assigned to the subfamily Deomyinae, which is closely related to the Gerbillinae. In another study, Chevret et al. (1993b) showed that Otomyinae are closely allied to the tribe Arvicanthini, thus unequivocally constituting a subset of the subfamily Murinae at the tribe level (Ducroz et al., 2001; Jansa and Weksler, 2004). Jansa and Weksler (2004) also strongly suggested that Lophiomyinae belonged to the Muridae. Only part of these proposals was followed by Musser and Carleton (2005) who recognized the following five subfamilies in the family Muridae: Deomyinae, Gerbillinae, Leimacomyinae, Murinae and Otomyinae. Nowadays the most consensual classification agrees on five subfamilies: Deomyinae (four genera and *ca.* 42 species), Gerbillinae (16 genera and *ca.* 103 species), Leimacomyinae (only one species, possibly extinct; Kingdon, 2015), Lophiomyinae (only one species) and Murinae (129 genera and *ca.* 584 species; see the review of Granjon and Montgelard, 2012).

**Table 1:** Brief summary of taxonomic changes in the family Muridae. The taxonomic position of particular lineages has changed significantly and they were either considered as separate families, or included in other families outside Muridae (names of families are in bold).

| Simpson (1945) |  | Chaline et al. (1977) |  | Lavocat (1978) |  |
| --- | --- | --- | --- | --- | --- |
| **Cricetidae** | Lophiomyinae | **Cricetidae** | Lophiomyinae | **Nesomyidae** | Lophiomyinae |
|  | Gerbillinae | **Gerbillidae** |  |  | Gerbillinae |
| **Muridae** | Murinae | **Muridae** | Murinae |  | Otomyinae |
|  | Otomyinae | **Nesomyidae** | Otomyinae | **Muridae** |  |
| Carleton and Musser (1984) |  | Musser and Carleton (2005) |  | Granjon and Montgelard (2012) |  |
| **Muridae** | Lophiomyinae | **Cricetidae** | Lophiomyinae | **Muridae** | Lophiomyinae |
|  | Gerbillinae | **Muridae** | Otomyinae |  | Gerbillinae |
|  | Murinae |  | Gerbillinae |  | Murinae |
|  | Otomyinae |  | Murinae |  | Deomyinae |
|  |  |  | Deomyinae |  | Leimacomyinae |
|  |  |  | Leimacomyinae |  |  |

Musser and Carleton’s (2005) comprehensive catalogue listed 730 species in the family Muridae. Estimates of species diversity in this family are very likely not definitive, as new murid taxa are being regularly described (e.g. Carleton et al., 2015; Esselstyn et al., 2015; Missoup et al., 2016; Mortelliti et al., 2016; Rowe et al., 2016a). Expected and ongoing raise in species number can be accounted for by an increased focus on poorly known regions with high levels of endemism, especially in tropical Asia and Africa. It is also linked with the development of integrative taxonomy studies, where molecular genetic approaches are able to detect taxa and geographical regions with high cryptic diversity (e.g. Bryja et al., 2014, 2017; Ndiaye et al., 2016).

Because of the high species richness of the family, determining the precise timing of its radiation is of particular paleobiogeographic interest. Several dated estimates for the age of Muridae are available owing to studies either focusing on the order Rodentia (Adkins et al., 2001, 2003; Fabre et al., 2012; Montgelard et al., 2008), on the superfamily Muroidea (Muroidea; Schenk et al., 2013; Steppan et al., 2004) or on various murid subsets (e.g. Bryja et al., 2014; Chevret and Dobigny, 2005; Dobigny et al., 2013; Fabre et al., 2013; Pagès et al., 2016; Rowe et al., 2008, 2011, 2016b). However, no clear consensus could be reached for the age of the family Muridae. Indeed, age estimates derived from all aforementioned studies are far from being congruent, likely because their datasets have not been designed for this particular purpose. In addition, all these studies used very diverse dating procedures, some of them relying on substitution rate calibrations (e.g. Arbogast et al., 2001; Nicolas et al., 2008) whereas others used fixed ages (e.g. the putative *Mus*/*Rattus* split at 12 Ma; Steppan et al., 2004), very distant fossil constrains (Adkins et al., 2003; Fabre et al., 2012; Montgelard et al., 2008) or primary calibrations using various fossil constraints within or outside the family Muridae (e.g. Bryja et al., 2014; Pagès et al., 2016; Rowe et al., 2016b; Schenk et al., 2013).

For fossil-based calibrations of molecular clocks, it is crucial: (i) to properly assign and place fossils on the tree, and (ii) to correctly estimate the age of fossil-bearing formations (Parham et al., 2012; Sauquet et al., 2012). Unfortunately the fossil record of oldest murids is quite fragmentary and mostly consists of isolated teeth and mandible remains, thus sometimes making taxonomic identification difficult. The earliest representatives for the family Muridae include the tribe Myocricetodontini with genera such as †*Myocricetodon* Lavocat, †*Dakkamys* Jaeger and †*Mellalomys* Jaeger (Jacobs and Flynn, 2005; Lazzari et al., 2011). Extinct members of the genus †*Potwarmus* Lindsey could be considered as a stem group of the subfamily Murinae based on detailed analyses of dental morphology (Lazzari et al., 2011; López Antoñanzas, 2009; Wessels, 2009). The earliest unequivocal representative of the subfamily Murinae is the genus †*Antemus* Jacobs (Jacobs and Downs, 1994; Jacobs and Flynn, 2005; Kimura et al., 2015). †*Antemus* possesses a new cusp (anterostyle, also known as t1), which is a synapomorphy of Murinae. The earliest record of †*Antemus chinjiensis* is dated at 13.8 Ma (Jacobs et al., 1990) based on specimens from the locality YGSP 491, Chinji Formation in the Potwar Plateau, Pakistan (Jacobs, 1977). In the fossil record of the Potwar Plateau, two more derived fossil genera are of particular interest: †*Karnimata* Jacobs and †*Progonomys* Schaub. Based on the relative position of the anterostyle to the lingual anterocone on M1, Jacobs (1978) hypothesized that †*Karnimata* is related to *Rattus* and that †*Progonomys* is a member of the lineage including *Mus*. Hence, their first stratigraphic occurrence has been used to define the widely used *Mus*/*Rattus* calibration (*ca.* 12 My; Jacobs and Downs, 1994). However, in 2015, Kimura et al. revisited these fossils from a paleontological perspective and proved this calibration point to be controversial. They showed that †*Karnimata* is a member of the Arvicanthini-Millardini-Otomyini clade rather than a member of the lineage encompassing the genus *Rattus* and its relatives (i.e. tribe Rattini). Therefore, they demonstrated that the continuous fossil record of the murine rodents from the Potwar Plateau actually provides a minimum age for the most recent common ancestor of the lineages leading to *Arvicanthis* Lesson and *Mus* (= *Mus*/*Arvicanthis* split).

Recent progresses in divergence dating analyses lead us to revisit results previously obtained by favouring the implementation of a new set of well-justified primary fossil calibrations within a Bayesian framework. In comparison to previous studies (listed above), our study can be considered as medium-sized in terms of taxonomic sampling, and essentially focused on the family Muridae. But our study stands out for rigorous evaluation of the fossil data for this highly diverse mammalian family. The present study has four main objectives: (i) to design a comprehensive multi-marker molecular dataset for the family Muridae, (ii) to review the murid fossil record in order to identify reliable and suitable primary fossil calibrations, (iii) to provide a reliable estimate of the timing of diversification of the family using multiple fossil calibrations, and (iv) to lean on the resulting dated phylogeny to reconstruct the biogeographic history of the family using up-to-date analytical approaches.

## 2. Material and Methods

### 2.1. Taxon sampling

For this study, new DNA sequences were generated for five murid species (*Acomys* cf. *cineraceus*, *Acomys subspinosus* (Waterhouse), *Acomys wilsoni* Thomas, *Arvicanthis niloticus* (Desmarest), *Arvicanthis neumanni* (Matschie) see Appendix A). Though we largely relied on GenBank data for this work, it is worth underlining that our research group generated thousands of murid sequences (all deposited in GenBank) in the past 15 years (we used some of these sequences for 44 species included in this study). In total, our dataset (Appendix A) encompasses 160 murid species representing 82 of the 151 known murid genera. All four extant subfamilies (if considering the Togo mouse from Leimacomyinae to be extinct) of Muridae are included. For the largest subfamily Murinae, we included representatives of all 10 tribes that have been defined by Lecompte et al. (2008): Apodemini, Arvicanthini, Hydromyini, Malacomyini, Millardini, Murini, Otomyini, Phloeomyini, Praomyini and Rattini. As outgroup taxa, we selected five species of the family Cricetidae (from subfamilies Arvicolinae, Cricetinae, Neotomyinae and Tylomyinae), which constitutes the sister group of Muridae (Fabre et al., 2012). Finally, the tree was rooted using *Calomyscus baluchi* Thomas, a representative of the more distant family Calomyscidae (Fabre et al., 2012). All species names followed Musser and Carleton (2005) and Monadjem et al. (2015).

### 2.2. DNA extraction, sequencing and molecular matrix

DNA was extracted using a Qiagen^®^ DNeasy Blood and Tissue kit (Qiagen, Hilden, Germany) following the manufacturer’s instructions. Two nuclear gene fragments were targeted using the following combinations of polymerase chain reactions (PCR) primers: IRBP217 and IRBP1531 (Stanhope et al., 1992) for the fragment of the ‘interphotoreceptor retinoid binding’ (IRBP) gene; RAG1F1705 and RAG1R2951 (Teeling et al., 2000) for a fragment of the ‘recombination activating gene 1’ (RAG1) gene. For PCR protocols, see Bryja et al. (2017) and Teeling et al. (2000), respectively. PCR products were Sanger sequenced in both directions using the BigDye^®^ Terminator chemistry (Thermo Fisher Scientific) either in the Institute of Vertebrate Biology on an ‘Applied Biosystems^®^ 3130xl Genetic Analyzer’, or commercially through the GATC Biotech company (Konstanz, Germany). New sequences were deposited in GenBank under accession numbers KY634246 to KY634250.

The newly generated sequences were further combined with data from GenBank. The resulting matrix (see Appendix A) encompasses the following six nuclear and nine mitochondrial gene fragments: ‘acid phosphatase 5’ (AP5), BRCA1, intronic portion of ‘Peripheral benzodiazapine receptor variant’ (BZRP), ‘growth hormone receptor’ (GHR), IRBP and RAG1, for the nuclear genes, and ‘12S ribosomal RNA’ (12S), ‘16S ribosomal RNA’ (16S), ‘ATP synthase 8’ (ATPase8), ‘cytochrome c oxidase I’ (COI), ‘cytochrome oxidase II’ (COII), ‘cytochrome *b*’ (Cytb), ‘Aspartic acid transfer RNA’ (tRNA-Asp), ‘Lysine transfer RNA’ (tRNA-Lys), ‘Serine transfer RNA’ (tRNA-Ser), for the mitochondrial genes. For nine taxa (*Acomys* cf. *cineraceus, Acomys wilsoni* Thomas, *Aethomys chrysophilus* (de Winton)*, Aethomys hindei* (Thomas)*, Aethomys kaiseri* (Noack)*, Aethomys silindensis* Roberts*, Arvicanthis nairobae* J.A. Allen*, Arvicanthis neumanni* and *Thallomys paedulcus* (Sundevall), gene fragments were concatenated from two individuals to minimize the amount of missing data. For all protein-coding genes, we used Mesquite 3.2 (Maddison and Maddison, 2007) to check the coding frame for possible errors or stop codons. The sequences of several markers (i.e. 12S, 16S, intronic portion of BZRP, tRNA-Asp, tRNA-Lys and tRNA-Ser) were variable in length; their alignment was accomplished using MUSCLE (Edgar, 2004) with default settings.

### 2.3. Phylogenetic analyses

Phylogenetic analyses were conducted using both Bayesian inference (BI) and maximum likelihood (ML). Analyses were performed on the online computer cluster CIPRES Science Gateway (Miller et al., 2010; www.phylo.org) and on the high performance computing (HPC) cluster hosted in the Centre de Biologie pour la Gestion des Populations (CBGP) in Montferrier-sur-Lez, France. For both phylogenetic analytical approaches, we carried out partitioned analyses to improve phylogenetic accuracy (Nylander et al., 2004). The molecular dataset was divided *a priori* into 33 partitions: we used three partitions for each of the protein-coding genes (AP5, ATPase8, BRCA1, COI, COII, Cytb, GHR, IRBP and RAG1) and one partition for each of the rRNA-tRNA genes (12S, 16S, tRNA-Asp, tRNA-Lys and tRNA-Ser) as well as the BZRP intronic portion. The best partitioning scheme and substitution models were determined with PartitionFinder 1.1.1 (Lanfear et al., 2014) using a greedy heuristic algorithm; because of the risk of over-parameterization associated with the high number of specified partitions, the ‘unlinked branch lengths’ option was chosen over the ‘linked branch lengths’ option. The Bayesian information criterion (BIC) was also preferentially used to compare partitioning schemes and substitution models following the recommendation of Ripplinger and Sullivan (2008).

PartitionFinder (based on BIC) identified the same three partitions for both BI and ML analyses: two partitions are associated with a Generalized-Time-Reversible (GTR +Γ +I) model and one partition is associated with a Hasegawa-Kishino-Yano (HKY +Γ +I) model (see Table 2).

**Table 2:**
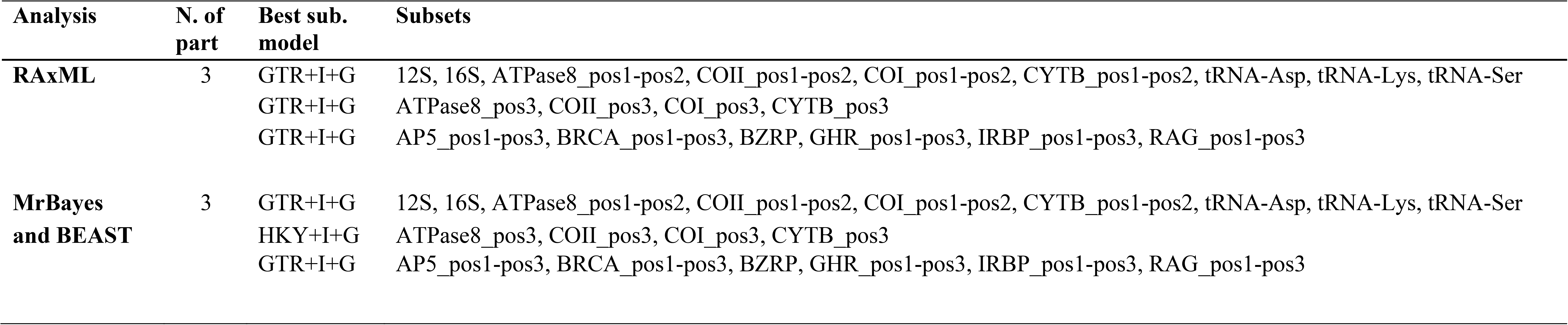
PartitionFinder results showing optimum partitioning schemes and best fit models for each analysis (RAxML, MrBayes, BEAST). Settings: BIC, unlinked branch lengths, greedy algorithm.

| Analysis | N. of part | Best sub. model | Subsets |
| --- | --- | --- | --- |
| RAxML | 3 | GTR+I+G | 12S, 16S, ATPase8_pos1-pos2, COII_pos1-pos2, COI_pos1-pos2, CYTB_pos1-pos2, tRNA-Asp, tRNA-Lys, tRNA-Ser |
|  |  | GTR+I+G | ATPase8_pos3, COII_pos3, COI_pos3, CYTB_pos3 |
|  |  | GTR+I+G | AP5_pos1-pos3, BRCA_pos1-pos3, BZRP, GHR_pos1-pos3, IRBP_pos1-pos3, RAG_pos1-pos3 |
| MrBayes and BEAST | 3 | GTR+I+G | 12S, 16S, ATPase8_pos1-pos2, COII_pos1-pos2, COI_pos1-pos2, CYTB_pos1-pos2, tRNA-Asp, tRNA-Lys, tRNA-Ser |
|  |  | HKY+I+G | ATPase8_pos3, COII_pos3, COI_pos3, CYTB_pos3 |
|  |  | GTR+I+G | AP5_pos1-pos3, BRCA_pos1-pos3, BZRP, GHR_pos1-pos3, IRBP_pos1-pos3, RAG_pos1-pos3 |

Bayesian inference analyses were carried out using MrBayes v3.2.6 (Ronquist et al., 2012b). Two independent runs with four MCMC (one cold and three incrementally heated chains) were conducted: they ran for 50 million generations, with trees sampled every 1,000 generations. A conservative 25% burn-in was applied after checking for stability on the log - likelihood curves and the split-frequencies of the runs. Support of nodes for MrBayes analyses was provided by clade posterior probabilities (PP) as directly estimated from the majority-rule consensus topology. Following Erixon et al. (2003), nodes supported by PP ≥ 0.95 were considered strongly supported.

Maximum likelihood analyses were performed using RAxML v8.2.8 (Stamatakis, 2014). Because this software does not allow simpler substitution models, we used three partitions with a General Time Reversible (GTR + Γ +I) model (see Table 2). The best ML tree was obtained using heuristic searches with 100 random addition replicates. Clade support was then assessed using a non-parametric bootstrap procedure with 1,000 replicates. Following Hillis and Bull (1993), nodes supported by bootstrap values (BV) ≥ 70 were considered strongly supported.

### 2.4. Evaluation of suitable fossil calibrations

Following the recommended criteria of Parham et al. (2012) for fossil calibrations, we rigorously compiled a list of potential candidates from the paleontological literature and eventually retained 18 candidate fossils (see Table 3). The candidate fossils possess the information for the collection site, unique identification number, and the state of preservation along with justification for the age of the fossil (i.e., age of fossil-bearing formation and stratigraphic level, preferably with an absolute age by radiometric dating and/or reliable relative age estimates, for example, by magnetostratigraphy).

**Table 3:**
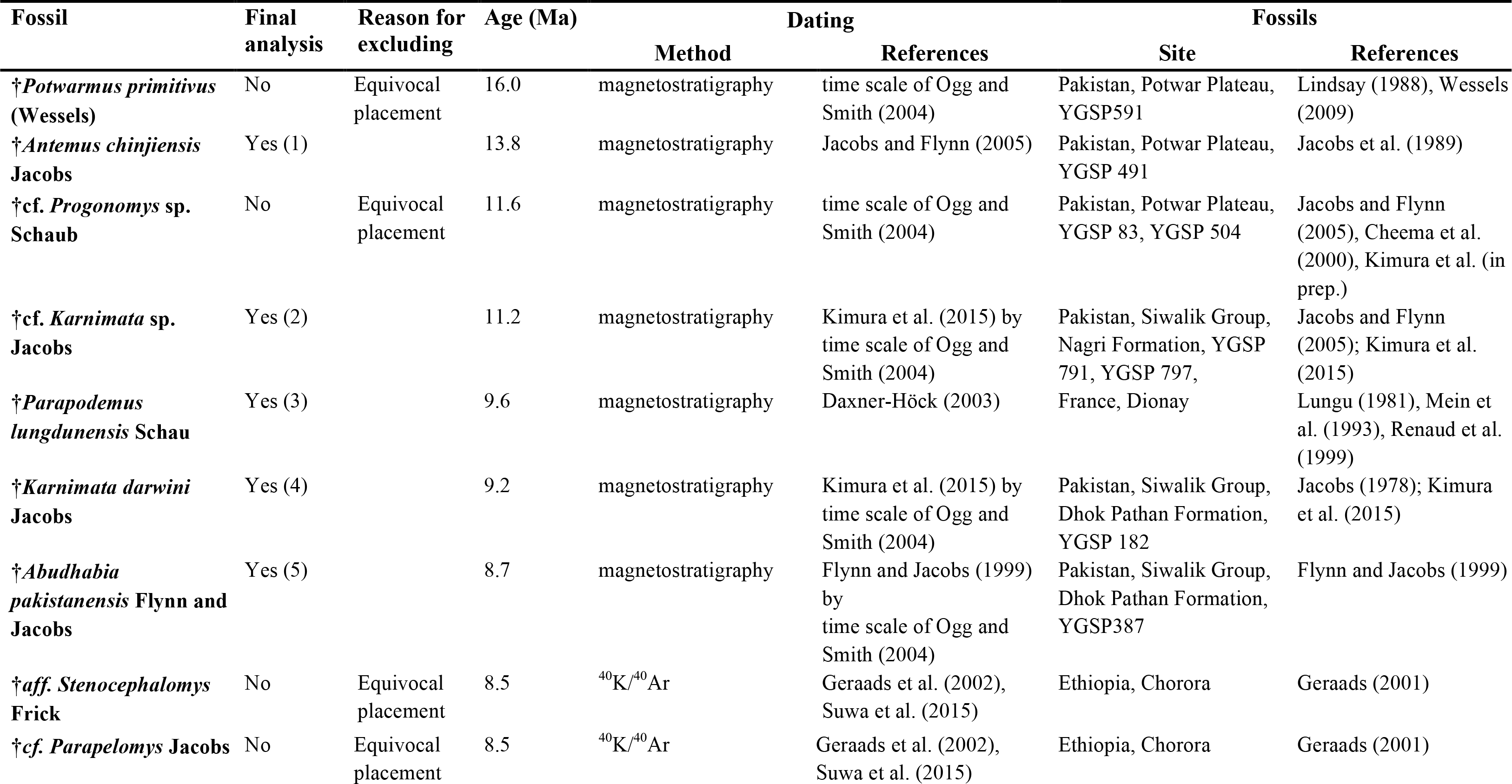
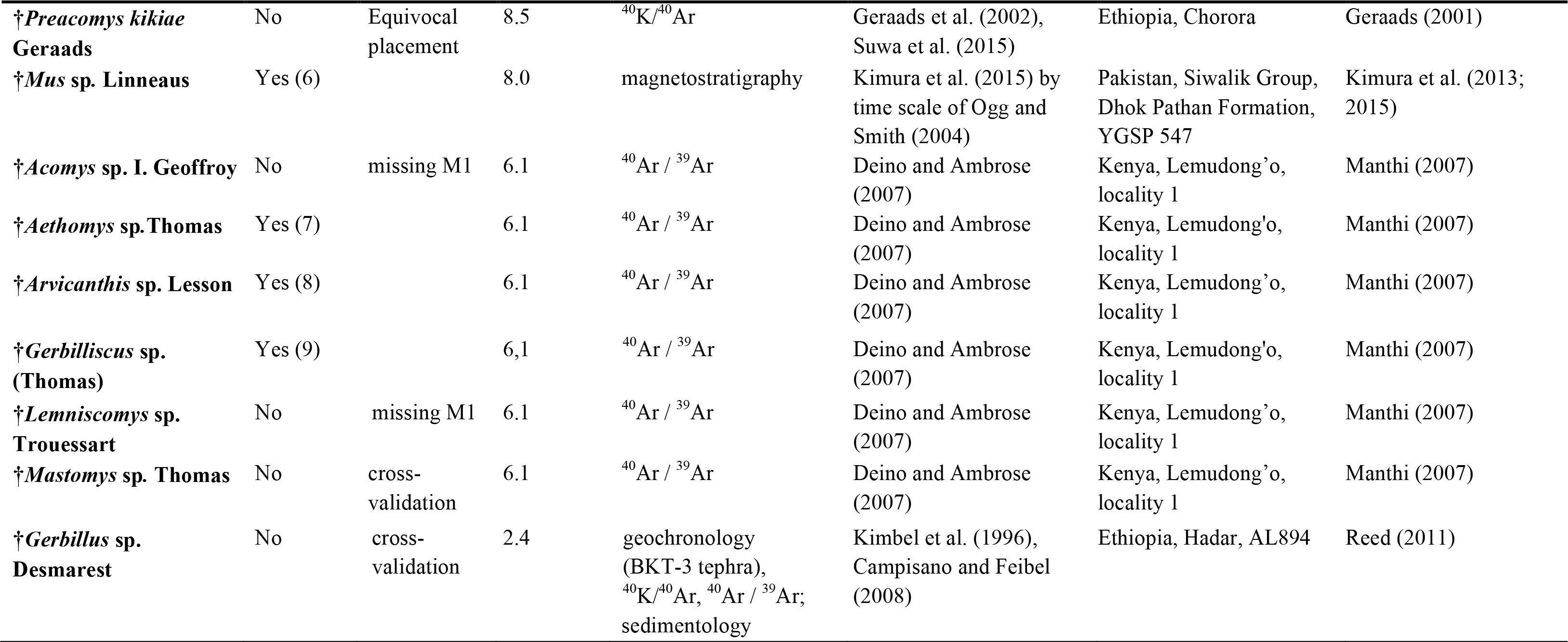
Overview of 18 candidate fossils with stratigraphic age, locality and relevant references. For more details see Appendix B.

In the next step, diagnostic morphological characters were reassessed to determine whether they could be reliably used as minimum age constraints in our phylogeny, either as crown or stem calibrations. Seven fossils were discarded following this step (see the ‘Results’ section).

For the remaining 11 fossils, we used the cross-validation procedure developed by Near and Sanderson (2004) and Near et al. (2005). The following approach was used: (i) we identified potential inconsistencies within the 11 remaining fossil calibrations, and (ii) we explored the impact of the inclusion of each of these fossils on our divergence time estimates. Each of the 11 fossil constraints was enforced at a time in a specific Bayesian relaxed-clock (BRC) analysis to estimate the ages of the remaining nodes (see also section 2.5). First, the sum of the squared differences between the molecular and fossil age estimates (SS) was calculated (for more details see Near and Sanderson, 2004). All calibration points were then ranked based on the magnitude of its SS score; here the fossil with the greatest SS score is assumed to be the most inconsistent with respect to all other fossils in the analysis (Near and Sanderson, 2004). Second, we calculated the average squared deviation, *s*, for all fossil calibrations in the analysis. Following the method of Near et al. (2005), we removed the fossil with the greatest SS score and recalculated *s* with the remaining fossil calibration points. This process was pursued until only the two fossil calibration points with the lowest and second lowest magnitudes of SS remained (Near and Sanderson, 2004). The rationale behind this procedure is to assess whether calibration points are approximately equally informative and accurate (Near et al., 2005): if it is the case the magnitude of *s* should only decrease by a small fraction whenever a fossil calibration is removed.

### 2.5. Bayesian relaxed-clock analyses

Although divergence time dating is now a well-established cornerstone of evolutionary biology, there is still no widely accepted objective methodology for converting data from the fossil record to calibration information of use in molecular phylogenies (Drummond and Bouckaert, 2015). In the last few years, several methodological approaches to better implement fossil calibrations have been developed, for instance allowing one to directly include fossil lineages in phylogenies (‘total-evidence dating’; Pyron, 2010; Ronquist et al. 2012a) or to account for information on the density of the fossil record (‘fossilized birth-death (FBD) process’; Stadler, 2010; Heath et al., 2014). However, for our study, a ‘total evidence dating’ approach was not applicable since it would have required the coding of a morphological matrix for both fossils and extant taxa, which is problematic given the fragmentary nature of muroid fossils. The use of the FBD methodology was also not envisioned because the fossil record of muroid rodents is too sparse. Instead, we relied on a node-dating approach in which fossil information is enforced on specific nodes through the use of parametric distributions.

Following our assessment of the murid fossil record and the results of cross-validation analyses, nine fossil calibrations were finally retained for the dating procedure (for more information, see Tables 3 and 4, and Appendix B). Five of them were defined based on fossil material collected in the Siwalik Group of Pakistan (†*Antemus chinjiensis*, †*Karnimata darwini* Jacobs, †cf. *Karnimata* sp., †*Mus* sp. and †*Abudhabia pakistanensis* Flynn and Jacobs). Three additional accepted fossils originate from 6.1 Ma fossils discovered in the Lemudong’o locality in Kenya (†*Aethomys* sp., †*Arvicanthis* sp. and †*Gerbilliscus* sp.), and the last retained fossil calibration constraint is defined by the 9.6 Ma fossil of †*Parapodemus lungdunensis* Schaub. Priors for fossil constraints were defined by using either uniform or lognormal statistical distributions in two separate analyses. Statistical distributions were bounded by the minimum ages provided by the fossil constraints and a conservative maximum age (*ca.* 25 Ma) for the root derived from the study of Schenk et al. (2013; see Table 4). In a preliminary way (see section 2.4 of the Material and Methods), BRC analyses were also conducted using one fossil constraint at a time to carry out cross-validation analyses.

**Table 4:** Overview of fossils finally selected for divergence dating, with parameters of uniform and lognormal prior distribution.

|  | Fossil | Position | Stem/Crown | Age (Ma) | Uniform distribution |  | Lognormal distribution |  |  |
| --- | --- | --- | --- | --- | --- | --- | --- | --- | --- |
|  |  |  |  |  | Min | Max | Offset | Log | Mean |
| 1 | †*Antemus chinjiensis* | crown Murinae | crown | 13.8 | 13.24 | 25.29 | 13.24 | 1.0 | 3.2 |
| 2 | †cf. *Karnimata* sp. | *Mus/Arvicanthis* split | crown | 11.2 | 10.47 | 25.37 | 10.47 | 1.0 | 4.0 |
| 3 | †*Parapodemus lugdunensis* | *Apodemus/Tokudaia* split | stem | 9.6 | 8.93 | 25.41 | 8.93 | 1.0 | 4.5 |
| 4 | †*Karnimata darwini* | TMRCA Millardini/Otomyini/Arvicanthini | crown | 9.2 | 8.52 | 25.42 | 8.52 | 1.0 | 4.6 |
| 5 | †*Abudhabia pakistanensis* | *Gerbilliscus/Desmodillus* split | crown | 8.7 | 8.01 | 25.43 | 8.01 | 1.0 | 4.7 |
| 6 | †*Mus* sp. | Murini | stem | 8.0 | 7.29 | 25.45 | 7.29 | 1.0 | 4.9 |
| 7 | †*Aethomys* sp. | *Aethomys* | stem | 6.1 | 5.34 | 25.50 | 5.34 | 1.0 | 5.5 |
| 8 | †*Arvicanthis* sp. | *Arvicanthis* | stem | 6.1 | 5.34 | 25.50 | 5.34 | 1.0 | 5.5 |
| 9 | †*Gerbilliscus* sp. | *Gerbilliscus* | stem | 6.1 | 5.34 | 25.50 | 5.34 | 1.0 | 5.5 |

Bayesian relaxed-clock analyses were conducted with BEAST v1.8.4 (Drummond et al., 2012) using uncorrelated lognormal relaxed clocks (Drummond et al., 2006). To limit the risk of over-parameterization: (i) we used three clock models (based on PartitionFinder results, Table 2); and (ii) we enforced a guide tree that corresponds to the topology with the best clade support (this topology corresponds to the topology obtained with MrBayes; see the ‘Results’ section). For the tree speciation model, a birth death process (Gernhard, 2008) was used in order to better account for extinct and missing lineages.

BEAST .xml files were modified to implement the path-sampling procedure for Bayes factor (B_F_) estimation following the recommendations of Baele et al. (2013). Out of the two calibrations, the calibration procedure with lognormal prior has the best harmonic mean (-208117.74 *versus* -208262.58 for the procedure with a uniform prior) and is recovered as the best-fit calibration procedure with a statistically significant BF of 289.68 (B_F_>10, Kass and Raftery, 1995). The final analysis (with nine verified fossil constraints and lognormal prior distribution for calibration constraints) was carried out by two independent runs each with 50 million generations and trees sampled every 5,000 generations. We used a conservative burn - in of 12.5 million generations per run. Post burn-in trees from both analyses were further combined using the LogCombiner module of BEAST. Convergence of runs was assessed graphically under Tracer v.1.6 and by examining the ESS of parameters.

### 2.6. Historical biogeography

Ancestral biogeography was reconstructed using the R package ‘BioGeoBEARS’ (Matzke, 2013). Data for species’ ranges were obtained from the International Union for Conservation of Nature website (https://www.iucn.org/). Five major biogeographic areas were defined on the basis of Olson et al. (2001): A, West Palearctic (from Western Europe to the Ural Mountains, including North Africa); B, East Palearctic (from the Urals to Japan); C, Indomalaya (from Afghanistan through the Indian subcontinent and Southeast Asia to lowland southern China, and through Indonesia as far as Java, Bali, and Borneo, west of the Wallace line); D, Australasia (Australia, New-Zealand, Papua-New-Guinea and neighbouring small islands); E, Afrotropics (Africa, northern part excluded). Dispersal rate between adjacent areas was fixed to 1 (A-B; B-C), whereas the dispersal of 0.7 (A-E; C-D) and 0.3 (B-D; B-E) was specified for long-distance dispersal or whenever a geographical barrier had to be crossed. Dispersal was disallowed between geographical areas separated by two or more areas (A-D; D-E). Six models of geographic range evolution were compared in a likelihood framework: (i) Dispersal-Extinction Cladogenesis model (DEC) similar to Lagrange (Ree and Smith, 2008), which parameterizes dispersal and extinction; (ii) DEC +J model (Matzke, 2013; 2014), which adds founder-event speciation with long-distance dispersal (cladogenesis, where daughter lineage is allowed to jump to a new range outside the range of the ancestor; Matzke, 2013) to the DEC framework; (iii) Dispersal Vicariance Analysis (DIVA; Ronquist, 1997); (iv) DIVA with long-distance dispersal (DIVA +J; Matzke, 2013); (v) Bayesian inference of historical biogeography for discrete areas (BayArea; Landis et al., 2013); and (vi) BayArea with long-distance dispersal (BayArea +J; Matzke, 2013). Model fit was assessed using the Akaike information criterion (AIC) and likelihood-ratio tests (LRT).

## 3. Results

### 3.1. Phylogeny of Muridae

Our multilocus dataset representing all major lineages of the family Muridae is 10,482 bp long with 42.5% missing data. Both BI and ML analyses yield similar topologies (see Fig. 1 for the topology inferred under BI, and Fig. S1 in Appendix D for the best-fit ML tree), as indicated by a high proportion of shared nodes (160 out of 162). BI and ML analyses differ only in the position of *Pelomys fallax* (Peters) and *Zyzomys argurus* (Thomas), but their placements are not significantly supported in either analysis. Clade support is moderate to high on average; if considering the number of nodes that are supported by PP ≥ 0.95 or BV ≥ 70%, BI analyses yield a slightly more robust topology (135 well-supported nodes) compared to the ML tree (122 well-supported nodes).

**Figure 1.**
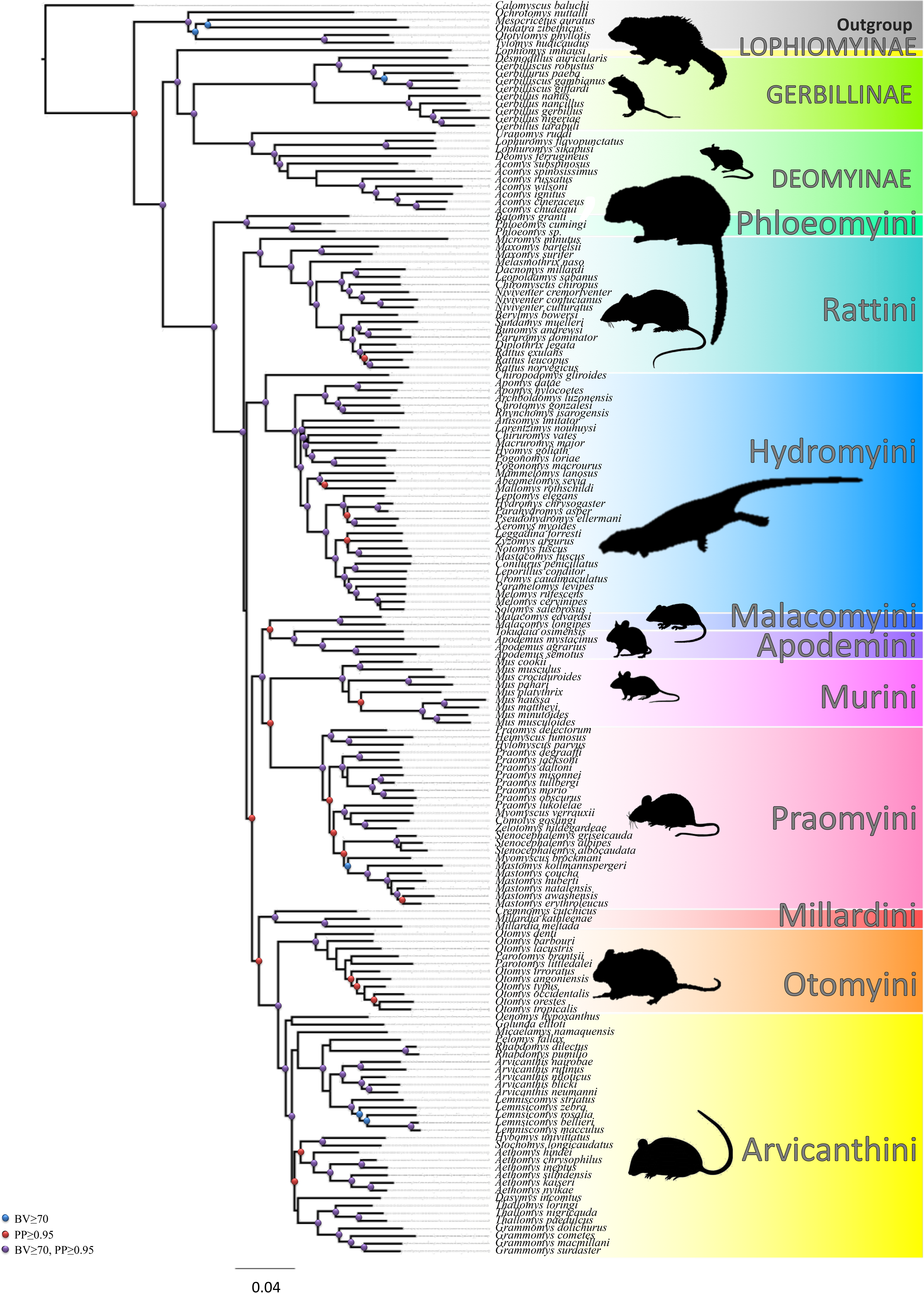
Molecular phylogeny of family Muridae based on Bayesian inference (BI) in MrBayes. Very similar topology was obtained also by maximum likelihood (ML) analysis in RAxML (see Appendix D). Red points show nodes supported in BI analysis (posterior probability PP ≥ 0.95), blue points show high bootstrap support in ML analysis (bootstrap BV ≥ 0.70). Violet nodes are supported by both analyses.

Phylogenetic analyses confirm the monophyly of the family Muridae, of all its four constituent subfamilies, as well as of the previously defined tribes of the subfamily Murinae (Fig. 1). On the contrary, the phylogenetic position of some genera (e.g. *Acomys* I. Geoffroy, *Dasymys* Peters, *Golunda* Gray, *Melomys* Thomas, *Micaelamys* Ellerman, *Pelomys* Peters, *Oenomys* Thomas and *Otomys* F. Cuvier) within particular tribes was only partly supported.

### 3.2. Evaluation of suitable fossil calibrations

We summarized all fossils considered in this study in Table 3 and Appendix B regarding taxonomic information and specification for prior settings (see also Figure 2 for their respective positions within the tree). Five out of 18 preselected fossils (i.e. †*Parapelomys robertsi* Jacobs, †*Potwarmus primitivus*, †*Preacomys kikiae* Geraads, †cf. *Progonomys* sp. Schaub and †aff. *Stenocephalomys* Frick) were excluded from further analyses because the scarcity of paleontological interpretation about their phylogenetic relationships impeded assigning them to specific nodes of the phylogeny (Appendix B). We also excluded fossils of *Acomys* and *Lemniscomys* Trouessart from the Lemudong’O locality, Kenya (Manthi, 2007), because first upper molars, which possess the most diagnostic characters in the murine dentition, are not described from the locality (Table 3; see more details also in Appendix B). The two-step cross-validation procedure resulted in a further reduction of the fossil set of possible calibration points. Specifically, we excluded two fossils: one is a 2.4 Ma fossil identified as the genus *Gerbillus* Desmarest, while the other one corresponds to a 6.1 Ma fossil identified as the genus *Mastomys* Thomas (Appendix C). The rationale is that (i) these two fossil calibrations exhibited the largest magnitude of SS, and that (ii) their removal also resulted in a very high (fivefold) decrease in *s* (see Appendix C for more details). As a result of the latter series of selection steps, nine fossils were finally retained for divergence dating (see Figure 2 for their position on the tree, Table 4 for specification of priors, and Appendix B for more details on all considered fossils).

**Figure 2.**
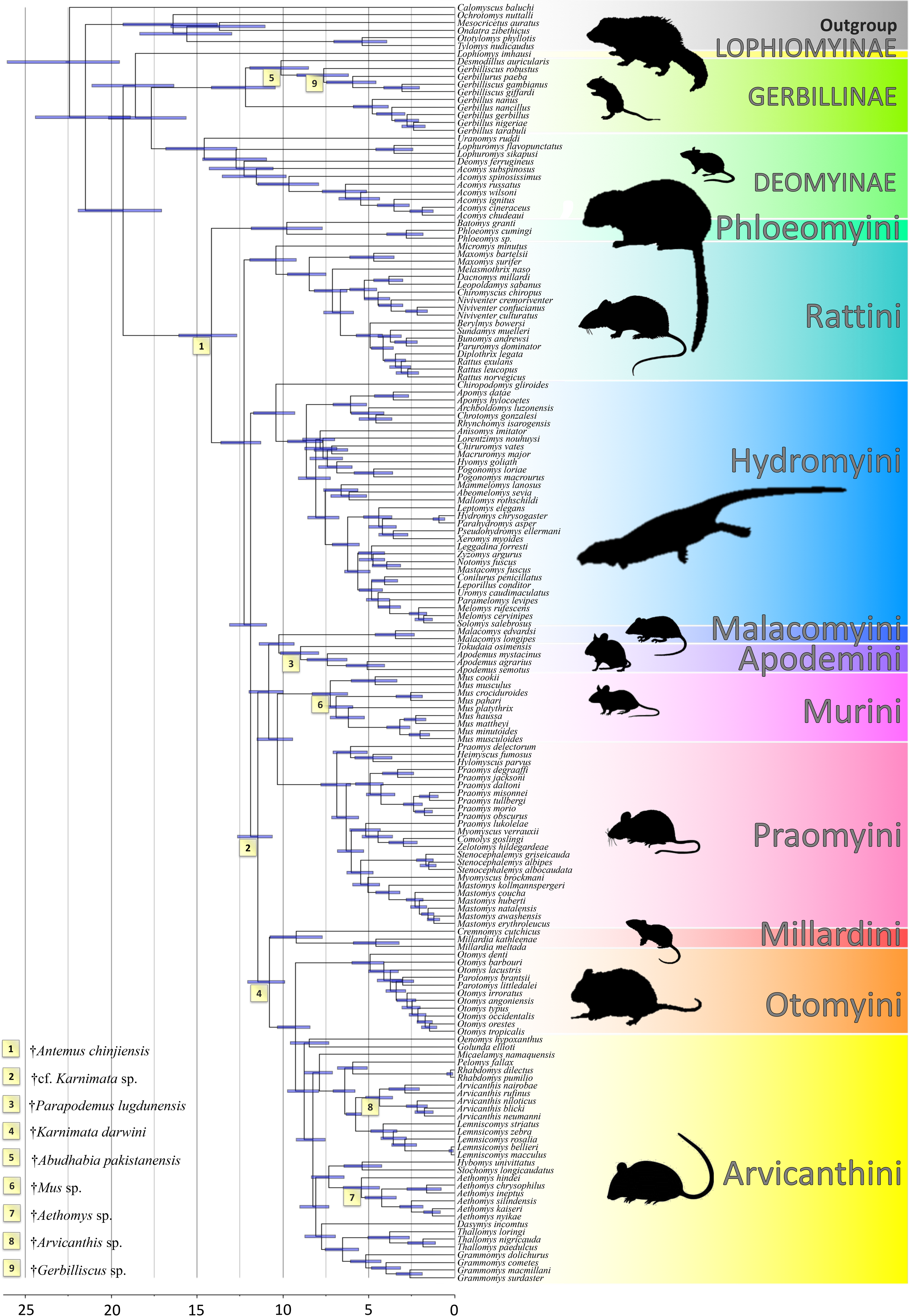
Divergence dating analysis of family Muridae. Nodes show medians of times to most recent common ancestor (MRCA), node bars indicate 95% HPD intervals. Latin numbers in yellow squares indicate positions of fossil constrains selected by multiple-step evaluation and used for final analysis (see Table 4 for more details).

### 3.3 Historical biogeography and divergence dating

Among the six models of geographic-range evolution compared in a likelihood framework in BioGeoBEARS, the Dispersal-Extinction Cladogenesis model with founder-event speciation (DEC +J) was chosen because of its best likelihood and AICc associated scores (lnL=-117.1, AICc=240.4; Table 5).

**Table 5:** Comparison of models used for BioGeoBEARS; likelihood scores (LnL), number of parameters (numparams), dispersal rate (d), extinction rate (e), free parameter controlling the relative probability of founder-event speciation events at cladogenesis (j), corrected Akaike Information Criterion (AICc), and AICc model weights.

|  | **LnL** | **numparams** | **d** | **e** | **j** | **AICc** | **AICc_wt** |
| --- | --- | --- | --- | --- | --- | --- | --- |
| DEC | -133.724 | 2 | 0.005 | 0.000 | 0.000 | 271.525 | 0.05181 |
| DEC +J | -129.826 | 3 | 0.004 | 0.000 | 0.008 | 265.806 | 0.90415 |
| DIVALIKE | -146.958 | 2 | 0.007 | 0.000 | 0.000 | 297.993 | 9.2665e-08 |
| DIVALIKE +J | -132.856 | 3 | 0.004 | 0.000 | 0.012 | 271.866 | 0.04369 |
| BAYAREALIKE | -178.311 | 2 | 0.005 | 0.036 | 0.000 | 360.698 | 2.2423e-21 |
| BAYAREALIKE +J | -137.687 | 3 | 0.003 | 0.000 | 0.014 | 281.528 | 0.00035 |

The dated tree resulting from the BRC analyses is shown in Figure 2 while dating estimates for all internal nodes are provided in Table 6, and results of ancestral distribution reconstructions are presented in Figure 3. The most recent common ancestor (MRCA) of Muridae originated during the early Miocene (median age of 19.3 Ma; 95% highest posterior density (HPD): 17.06-21.92 Ma) in the Afrotropical and Indomalayan bioregions.

**Figure 3.**
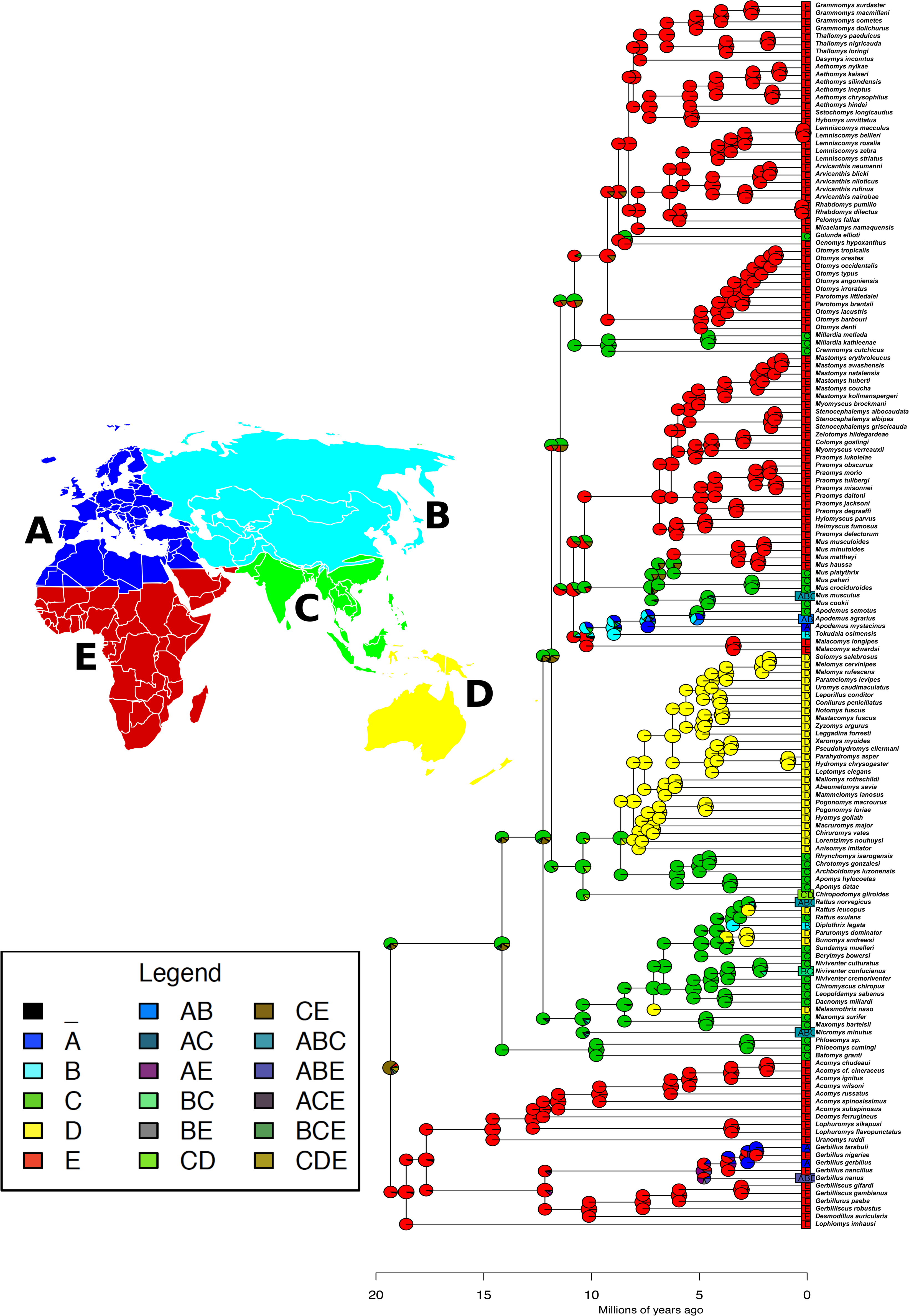
Ancestral reconstruction for family Muridae with BioGeoBEARS (DEC+J; d=0.008; e=0; j=0.0246; LnL=-117.11). Five biogeographical areas are represented using different colours: A, West Palearctic (dark blue); B, East Palearctic (light blue); C, Indomalaya (green); D, Australasia (yellow); E, Afrotropics (red).

**Table 6:** Results of divergence dating analysis. Time to the most common ancestor (MRCA) is shown as median in Ma, with 95% highest posterior density (HPD). Estimates from previous studies dealing with divergence dating of murid rodents are reviewed here for comparison.

| **TMRCA** | **median** | **2.5%** | **97.5%** | **Steppan et al.**<br>**(2004)** | **Lecompte et al.**<br>**(2008)** | **Rowe et al.**<br>**(2011)** | **Fabre et al.**<br>**(2012)** | **Fabre et al.**<br>**(2013)** | **Schenk et al.**<br>**(2013)** | **Bryja et al.**<br>**(2014)** | **Rowe et al.**<br>**(2016b)** |
| --- | --- | --- | --- | --- | --- | --- | --- | --- | --- | --- | --- |
| Muridae | 19.3 | 17.06 | 21.92 | 22.5 |  |  | ~33-23 |  | ~21.0 |  |  |
| Lophiomyinae* | 18.6 | 16.35 | 21.11 |  |  |  |  |  |  |  |  |
| Gerbillinae | 12.2 | 10.46 | 14.16 | 9.3 |  |  | ~23-5 |  | ~10.0 |  |  |
| Deomyinae | 14.6 | 12.68 | 16.82 | 13.1 |  |  | ~23-5 |  | ~13.0 |  |  |
| Murinae | 14.2 | 12.70 | 16.07 | 12.0 | 12.3 | 13.3 | ~23-5 |  | ~14.5 |  | ~14.0 |
| "core Murinae" | 12.3 | 11.28 | 13.62 | 10.3 | 11.3 |  |  | 11.60 | ~12.5 |  | ~12.5 |
| Phloeomyini | 9.8 | 7.72 | 11.85 |  | 8.6 |  |  |  | ~10.0 |  | ~10.5 |
| Rattini | 10.4 | 9.23 | 11.93 |  | 9.7 |  |  | 8.70 | ~11.0 |  | ~11.0 |
| Hydromyini | 10.4 | 9.31 | 11.71 |  | 8.9 |  |  |  | ~11.0 |  | ~11.0 |
| Malacomyini | 3.4 | 2.35 | 4.60 |  |  |  |  |  |  | 4.4 |  |
| Apodemini | 9.0 | 7.92 | 10.13 |  | 9.6 |  |  |  | ~7.5 | 9.5 | ~7.5 |
| Murini | 7.2 | 6.24 | 8.29 |  | 6.6 | 5.3 |  |  | ~6.0 | 7.4 | ~6.5 |
| Praomyini | 6.8 | 6.05 | 7.77 |  | 7.6 |  |  |  | ~5.0 | 6.8 | ~6.0 |
| Millardini | 9.2 | 7.72 | 10.75 |  |  |  |  |  |  |  |  |
| Otomyini | 4.9 | 4.12 | 5.98 |  |  |  |  |  | ~6.0 | 3.8 | ~3.0 |
| Arvicanthini | 8.8 | 7.92 | 9.72 |  | 8.4 | 7.3 |  |  | ~7.5 | 7.8 | ~8.5 |

\* offshoot from Gerbillinae + Deomyinae

Three subfamilies (Deomyinae, Gerbillinae and Lophiomyinae) belonging to the same clade started their diversification in the Afrotropics. Within this clade, a first split occurred *ca*. 18.6 Ma (95% HPD: 16.35-21.11 Ma) between the Lophiomyinae and the clade encompassing the Deomyinae and Gerbillinae. Deomyinae started their diversification *ca*. 14.6 Ma (95% HPD: 12.68-16.82 Ma) while Gerbillinae started theirs *ca*. 12.2 Ma (95% HPD: 10.46-14.16 Ma). In Deomyinae and Gerbillinae, several lineages were able to colonize the Palearctic region from the Afrotropics (in our dataset, this concerns *Acomys russatus* (Wagner) and *Gerbillus gerbillus* Olivier).

The subfamily Murinae originated in the Indomalayan region during the middle Miocene (median age of 14.2 Ma; 95% HPD: 12.70-16.07 Ma); the corresponding basal split separated the Phloeomyini and all remaining murines (‘core murines’ *sensu* Steppan et al. 2005). The next major split occurred in the Indomalaya between Rattini and the remaining tribes of Murinae (median age of 12.3 Ma; 95% HPD: 11.28-13.62 Ma), with an origin of Rattini estimated at 10.4 Ma (95% HPD: 9.23-11.93 Ma). The ancestral area of Rattini was also inferred to be the Indomalaya; during the course of their diversification, a few taxa colonized Australasia (e.g. *Bunomys andrewsi* (J.A. Allen), *Melasmothrix naso* Miller and Hollister, *Paruromys dominator* Thomas and *Rattus leucopus* (Gray)) as well as the West and East Palearctic (e.g. *Micromys minutus* (Pallas) and *Diplothrix legata* (Thomas)). The Hydromyini split from the remaining Murinae at *ca.* 11.9 Ma (95% HPD: 10.95-13.10 Ma); although basal lineages of this tribe are currently found in the Indomalaya (e.g. *Archboldomys luzonenzis* Musser, *Apomys datae* (Meyer), *Apomys hylocoetes* Mearns, *Chrotomys gonzalesi* Rickart and Heaney, *Chiropodomys gliroides* (Blyth) and *Rhynchomys isarogensis* Musser and Freeman), a specific and diverse lineage of Hydromyini also colonized and radiated in the Australasia *ca.* 8.1 Ma (95% HPD: 7.22-9.09 Ma). The clade gathering Apodemini, Malacomyini, Murini and Praomyini likely originated in the Afrotropics, with several lineages secondarily colonizing the Indomalaya and the West and East Palearctic. The split between Malacomyini (which remained in the Afrotropics) and Apodemini (which dispersed and differentiated mainly in the West and East Palaearctic) is estimated at *ca.* 10.2 Ma (95% HPD: 9.33-11.39 Ma). Murini started to diversify in the Indomalaya at *ca.* 7.2 Ma (95% HPD: 6.24-9.29 Ma). The intense radiation (51 extant species, Monadjem et al., 2015) of Praomyini occurred in the Afrotropics (median age of 6.8 Ma for the MRCA of Praomyini; 95% HPD: 6.06-7.77 Ma). The Indomalayan Millardini split from the predominantly Afrotropical Arvicanthini + Otomyini tribes at *ca.* 10.8 Ma (95% HPD: 9.88-12.04 Ma). The respective first diversifications within Arvicanthini, Millardini and Otomyini are estimated at *ca.* 8.8 Ma (95% HPD: 7.92-9.72 Ma), 9.2 Ma (95% HPD: 7.72-10.75 Ma) and 4.9 Ma (95% HPD: 4.12-5.98 Ma), respectively. The position of Asian *Golunda* within Arvicanthini is not resolved (Fig. 1); the dispersal to Indomalaya of the lineage leading to the extant *Golunda* species at *ca.* 8.5 Ma (95% HPD: 7.33-9.57 Ma; as suggested in Fig. 3) should therefore be taken with caution.

## 4. Discussion

### 4.1. Selection of taxa and molecular markers

Our sampling of 160 species from 82 genera represents 22% of known murid species diversity and more than half of the generic diversity of the family Muridae. When one compares our sampling effort to previous studies (Table 7), only the study of Fabre et al. (2012) relied on a better sampling for the family Muridae (302 species from 105 genera, i.e. about 41% of known species diversity). In the study of Schenk et al. (2013), 18% of murid species are included. The number of sampled murid species is also lower in Lecompte et al. (2008) and Rowe et al. (2008) because their studies focussed on specific tribes and subfamilies.

**Table 7:** Comparison between previous relevant studies.

| Author | Focus | Subfamilies | # genera | # species | Fossils in Muridae | # genetic markers |
| --- | --- | --- | --- | --- | --- | --- |
| Fabre et al. (2012) | Rodentia | 4 | 105 | 302 | 0 | 11 (8)* |
| Schenk et al. (2013) | Muroidea | 4 | 85 | 136 | 4 | 4 |
| our study | Muridae | 4 | 82 | 160 | 9 | 15 |
| Rowe et al. (2008) | Hydromyini (outgroup all Murinae + Gerbillinae+Deomyinae) | 3 | 66 | 78 | 2 | 8 |
| Lecompte et al. (2008) | Murinae | 3 | 46 | 83 | 2 | 3 |

\* 11 genetic markers for Rodentia, only 8 for Muridae

### 4.2. Calibration of molecular clock and divergence dating

Using the classical *Mus/Rattus* calibration as prior for divergence dating often lead to an underestimation of the age of the subfamily Murinae, inferring median ages that are generally comprised between 13.3 to 12.0 Ma. Only the most recent studies (e.g. Rowe et al., 2016b) used the correct *Mus/Arvicanthis* calibration with a prior median age of 11.1 Ma (as suggested by Kimura et al., 2015). This resulted in the estimation of Murinae age of *ca*. 14.0 Ma (Rowe et al., 2016b), which is consistent with our study (Table 6).

Other fossils frequently used for molecular clock calibration are from the genus †*Parapodemus*. Recent studies used these fossils in two ways: the first occurrence of †*Parapodemus* sp. (Martín-Suáres and Mein, 1998) in the late Miocene in Europe (Lungu, 1981; see Appendix B) was used to calibrate the MRCA of ‘Apodemurini’ (representing the split between Apodemini+Malacomyini and Murini+Praomyini; Fabre et al., 2013; Rowe et al., 2011). Another calibration point is based on the discovery of †*Parapodemus pasquierae* Aguilar and Michaux, from ‘Lo Fournas 6’ site (Roussillon, France). Authors postulated that the latter species and the smaller †*Parapodemus lugdunensis* co-occurred during the same time period ‘MN10’ (Aguilar et al., 1999; Montuire et al., 2006), dated approximately at 9.7 Ma (Mein, 2003). Michaux et al. (2002) considered the differences between these two species as representative of the split between the large *Apodemus mystacinus* Danford and Alston and all smaller *Apodemus* species from the subgenus *Sylvaemus*, but they used a younger age of 7.0 Ma as a prior for their divergence. Numerous authors followed this calibration (e.g. Bryja et al., 2014; Fabre et al., 2013; Lecompte et al., 2008; Schenk et al., 2013) even if there is no clear rationale for it. The estimated dates of MRCA of Apodemini range from 7.5 Ma (Rowe et al. 2016b; Schenk et al., 2013) to 9.6 Ma (Bryja et al., 2014; Michaux et al., 2002; Lecompte et al., 2008). In our study, we conservatively used the †*Parapodemus lugdunensis* fossil as a stem constraint for Apodemini and this placement resulted in an estimation of their MRCA at 9.0 Ma (95% HPD: 7.92-10.13 Ma).

There are several localities in Africa (e.g. Lukeino Formation, Winkler, 2002; Lemudong’o, Ambrose et al., 2007; Manthi, 2007) where fossil representatives of *Arvicanthis* were identified. These fossils were used for molecular clock calibration in several studies with a prior MRCA for the genus at 6.1 Ma (e.g. Fabre et al., 2013; Rowe et al., 2011). The fossils from Lemudong’o were also used, in a less conservative way, by Bryja et al. (2014): based on early records of †*Otomys* sp. (*ca.* 5.0 Ma; Denys, 1990), †*Aethomys* sp., †*Arvicanthis* sp., †*Lemniscomys* sp. (from Lemudong’o = 6.1 Ma; Ambrose et al., 2007; Manthi et al., 2007) and other relevant samples where these and related genera were absent (9.50-10.50 Ma; Mein et al. 2004), they set the split between Arvicanthini and Otomyini within 6.08-9.54 Ma. This calibration resulted in an estimation of the MRCA for the tribe Arvicanthini at 7.8 Ma (Bryja et al., 2014), which is about 1 million year younger than our own estimate (Table 6). Rowe et al. (2016b) used as minimum age 8.7 Ma for the split between *Arvicanthis* and *Otomys* based on the study of Kimura et al. (2015) that set minimum and maximum ages for locality Y388, where †*Karnimata darwini* was found. This calibration resulted in an estimated MRCA of Arvicanthini at 8.5 Ma. In our study, we instead used the age of an older locality (Y182; median age of 9.2 Ma) where †*Karnimata darwini* was also found (Jacobs, 1978; Kimura et al., 2015), in order to set a crown calibration for the Arvicanthini/Millardini/Otomyini clade. This placement resulted in an estimation of their MRCA at 8.8 Ma (95% HPD: 7.92-9.72 Ma) (Table 6).

### 4.3. Historical biogeography with focus on faunal exchanges between the Afrotropics and the Indomalaya

Our study could not resolve the origin of murid rodents, but it was either in the Afrotropics or in the Indomalaya. Our inferred ancestral tropical range for the MRCA of murids is consistent with the fact that most extant murid taxa are still distributed in warm and moist tropical areas. During the Early Miocene (23.0-16.0 Ma), the rotation of Africa and Arabia, and finally the collision with Eurasia formed a landbridge between Africa and Eurasia (the so-called ‘*Gomphotherium* landbridge’; Rögl, 1999). During this time period, early murids colonized both geographical regions. The subsequent reopening of the Mediterranean-Indo-Pacific seaway (‘Indo-Pacific recurrence’; Rögl, 1999) separated Africa from Eurasia again, thus giving rise to the main clades of Afrotropics and Indomalaya rodents. Three subfamilies, Deomyinae, Gerbillinae and Lophiomyinae, then likely diversified in the Afrotropics (Chevret and Dobigny, 2005; Ndiaye et al., 2016; Schenk et al., 2013, this study; Figure 3). This hypothesis is supported by paleontological records since the oldest fossils tied to these subfamilies were found in the Afrotropics (e.g. late Miocene *Acomys, Gerbilliscus, Lophiomys* and †*Preacomys* from East Africa; Winkler et al. 2010 and references therein). The subfamily Murinae started to diversify in Indomalaya, most probably in Southeast Asia, where we can also find the hitherto highest phylogenetic diversity, including the oldest offshoots of this clade (e.g. the ancestor of Phloeomyini probably lived in the Philippines, those of Rattini and Hydromyini in South-east Asia, etc.; Fabre et al., 2013).

During the Middle Miocene (16.0-11.6 Ma), the Mediterranean-Indo-Pacific seaway closed again at the beginning of the Serravallian *ca.* 13.8 Ma (‘Parathethys Salinity Crisis’; Rögl, 1999), co-incidentally with a global cooling that caused vegetation shifts and a general aridification (Prista et al., 2015). The newly formed landbridge (Rögl, 1999) allowed repeated dispersals of murine rodents from Asia to both Africa and Eurasia. Murine fossil records provide clear evidence for connections between the Indomalaya, the Palearctic, and the Afrotropics. Among them, there are two conspicuous examples: (i) †*Progonomys* was recorded in many Indomalayan Middle Miocene localities (Jacobs and Flynn, 2005) as well as in the Palearctic region (Algeria: Wessels, 2009; China: Qiu et al., 2004; Egypt: Heissig, 1982; France: Mein et al., 1993; Spain: Weerd, 1976); and (ii) the oldest records of †*Parapelomys* spp. were found synchronously in Africa (8.5 Ma; Chorora, Ethiopia; Geraads, 2001) and in Pakistan (*ca.* 8.0 Ma; Jacobs and Flynn, 2005). During this period, representatives of several murine tribes occurred in the Afrotropics (Arvicanthini, Malacomyini, Otomyini and Praomyini) and the Indomalaya (Millardini, Murini, Rattini, and basal lineages of Hydromyini).

The last faunal interchange of murid taxa between Africa, Asia and Western Palearctic (Benammi et al., 1996; Sabatier, 1982; Sen, 1977, 1983; Winkler, 2002) is coincident with Messinian Salinity Crisis *ca.* 6 Ma during the Late Miocene (Hsü et al., 1973, 1978). During this period of global sea level depression (Haq et al., 1987) Africa and Arabia were reconnected through Neguev-Sinai landbridge (‘Levantine corridor’, Fernandes et al., 2006) and landbridge in the Bab-el-Mandeb (Bosworth et al., 2005). In murids, evidence to support this faunal exchange can be found in the African subgenus *Nannomys* (genus *Mus*), which colonized Afrotropics and started there its radiation *ca.* 5.2 Ma (Bryja et al., 2014). A possible example for an opposite west-to-east migration is the genus *Golunda*, which belongs to the Arvicanthini tribe. In a predominantly Afrotropical clade, *Golunda* is the only genus that occurs in the Indomalaya, probably since the end of Miocene (Ducroz et al., 2001; Fig. 3). However, one should be cautious with this scenario since the position of *Golunda* within Arvicanthini is not well supported (Fig. 1). Africa-to-Asia dispersals at the Miocene/Pliocene boundary have been also recorded in other taxa, such as rodents (e.g. *Myomyscus yemeni* (Sanborn and Hoogstraal); our unpubl. data), reptiles (e.g. *Varanus yemenensis*: Böhme et al., 2003, Portik and Papenfuss 2012; *Hemidactylus* geckos: Šmíd et al. 2013; *Echis* vipers: Pook et al. 2009) and hamadryas baboons (Winney et al. 2004).

## 5. Conclusion and perspectives

In this study, we provided an improved multilocus dated phylogeny for the highly speciose family Muridae. Both our dating and historical biogeography analyses suggest that the family originated during the Early Miocene, and subsequently gave rise to four extant subfamilies: three in the Afrotropical region (Deomyinae, Gerbillinae and Lophiomyinae) and one in the Indomalaya (Murinae). Our study also supports a dynamic biogeographic scenario in which repeated colonisation events occurred in the Australasian (Hydromyini, Rattini), Afrotropical (Malacomyini, Praomyini, Arvicanthini, Otomyini) and Palearctic (Apodemini) regions. One of the strong aspects of this study lies in the assessment and treatment of fossil data (Appendix B); such data is likely to be useful for further studies investigating the timing of diversification of rodents, or even mammals in general. For an easy access to all corresponding fossil records, we have made data available on the Date-a-Clade Website (http://palaeo.gly.bris.ac.uk/fossilrecord2/dateaclade/index.html), Paleobiology Database (http://fossilworks.org/) and TimeTree Database (http://timetree.org).

## Acknowledgments

This study was supported by two projects of the Czech Science Foundation, no. 15-20229S, the Ministry of Culture of the Czech Republic (DKRVO 2017/15, National Museum, 00023272) and JSPS KAKENHI JP15H06884 (Grant-in-Aid for Young Scientists Start-up). Most analyses were run on CBGP HPC computational platform, while a minor part was performed on CIPRES Gateway. We thank Alexandre Dehne Garcia for his help on the HPC CBGP cluster and Nick J. Matzke for introduction into BioGeoBEARS analysis. We would also like to thank Alisa Winkler and Christiane Denys for discussion about some paleontological aspects, as well as Arame Ndiaye and Pascal Chevret for providing additional sequences. We are grateful to Vincent Lazzari, Fredrick K. Manthi, Pierre Mein, Rajeev Pantanik and Sevket Sen who provided permission for the reproduction of the figures in Appendix B.

## Authors’ contributions

GJK, GD, LG, JB conceived the ideas; TA collected genetic data (including new genotyping); YK, TA collected and analysed paleontological data; TA, GJK performed phylogenetic analyses; TA, JB, GJK wrote the first version of the manuscript that was then implemented by all authors.

## Supplementary material

**Appendix A.** List of taxa and genetic markers.

**Appendix B.** Description of considered fossils.

**Appendix C.** Results of cross-validation of fossil constraints.

**Appendix D.** Maximum likelihood phylogenetic tree.

